# Mechanism of neuroinvasion and interferon response in human choroid plexus organoids during Japanese encephalitis virus infection

**DOI:** 10.64898/2025.12.03.692106

**Authors:** Yothin Kumsang, Atsadang Boonmee, Chatpong Pethrak, Pisut Pongchaikul, Usanarat Anurathapan, Chatchai Muanprasat, Phisit Khemawoot, Bunpote Siridechadilok, Natapong Jupatanakul, Nithi Asavapanumas, Nunya Chotiwan

## Abstract

Orthoflaviviruses are medically important arthropod–borne pathogens that cause a significant global disease burden. Some of these viruses can cause neurological infections, requiring a clear understanding of how they enter the central nervous system (CNS). The choroid plexus (ChP) is a cerebrospinal fluid (CSF)–producing tissue residing in the brain ventricles. The ChP forms a blood–CSF barrier, which acts as a crucial peripheral–CNS interface and a physical defense against neuroinvasive pathogens. While this tissue is increasingly recognized as a potential entry site for hematogenous pathogens, how orthoflaviviruses overcome the blood–CSF barrier and the corresponding intrinsic immune response in a human model remains critically undefined. This study aimed to investigate the potential of the ChP to serve as a neuroinvasive site for orthoflaviviruses. ChP organoids (ChPOs) derived from human inducible pluripotent stem cells were established and determined for susceptibility to a panel of orthoflaviviruses, namely Japanese encephalitis virus (JEV), Zika virus (ZIKV), and Dengue virus (DENV). These viruses exhibited differential susceptibility to ChPOs, with the pathogenic JEV strain being the most infective. Transcriptomic analysis of ChPOs confirmed a robust interferon (IFN) response to JEV infection. Functional studies further demonstrated that antiviral defenses are crucial for controlling virus infection, and that type I IFN responses are superior to the epithelial–restricted type III IFN responses. Critically, despite the productive release of JEV into the CSF–like fluid within the ChP lumen, selective barrier integrity was maintained, confirming a nondestructive transcellular neuroinvasion mechanism across the blood–CSF barrier. This study highlights a valuable *in vitro* platform and its potential for studying virus neuroinvasion at the blood–CSF barrier.

## Background

The genus *Orthoflavivirus* comprises a group of arthropod–borne, single–stranded, positive–sense RNA viruses belonging to the family Flaviviridae. Medically important members of this genus include Dengue virus (DENV), Zika virus (ZIKV), Yellow fever virus (YFV), West Nile virus (WNV), Tick–borne encephalitis virus (TBEV), and Japanese encephalitis virus (JEV). Hundreds of millions of orthoflavivirus–infected cases are reported annually, with transmission distributed globally but concentrated in tropical and subtropical areas [1]. JEV, which is endemic to the Asia Pacific and South Asia regions, is transmitted by *Culex* mosquitoes, and its transmission cycle is maintained in a zoonotic cycle via avian and porcine reservoir hosts [2]. Human JEV infections are accidental, and humans are dead–end hosts for the virus. Most JEV infections affect children and are asymptomatic or result in mild febrile illness. However, approximately 1 in 250 JEV cases develop severe neurological manifestations, including encephalitis, meningoencephalitis, and meningitis [3]. Up to 30% of these severe cases are fatal, and 30–50% of those who survive suffer permanent neurological deficits.

Despite the availability of effective vaccines, JEV remains a significant global health concern. The disease burden is estimated to cause 100,000 apparent cases and 25,000 deaths annually [4]. Furthermore, the geographic range of JEV is expanding to new areas, such as Australia and higher altitudes in Nepal and Tibet [5, 6]. This expansion, together with increased international travel, heightens the risk of exposure for nonimmunized individuals, making the study of JEV increasingly critical. In particular, a better understanding of the host–virus interaction at peripheral–CNS interfaces is needed for mitigating devastating neurological outcomes.

The initial JEV infection occurs locally in the immune cells in the skin or in local lymph nodes near the site of inoculation. After local replication, the virus can enter the bloodstream, causing viremia, and it can then disseminate systemically to infect other organs, including the central nervous system (CNS) [7]. In a healthy human, the CNS is protected from the passage of viruses and other infective or toxic agents in the bloodstream by specialized neurovascular tissues called brain barriers, such as the blood–brain barrier (BBB). However, compromise of these barriers can facilitate the unrestricted entry of these substances like JEV into the CNS. Previous *in vivo* studies on JEV–infected rodents have demonstrated JEV infiltration of peripheral immune cells, extensive neuroinflammation, and BBB leakage. However, these studies also reported the presence of JEV RNA in the brain prior to any significant BBB disruption event [8, 9], suggesting BBB leakage might be the effect of the infection within the CNS and the primary neuroinvasion mechanism remains unraveled, perhaps involving a brain barrier other than the BBB.

One potential brain barrier that could be compromised by JEV infection is the blood–cerebrospinal fluid (CSF) barrier, which is a constituent of a secretory tissue called the choroid plexus (ChP) [10]. The ChP, which is found within the lateral, third, and fourth ventricles of the brain, has a structure consisting of fenestrated capillaries (also known as leaky vessels) that are enveloped by connective tissue and a layer of epithelial cells. These epithelial cells, which are the main producers of CSF, form interconnected tight junctions and regulate the passage of substances from the peripheral circulation into the CSF. Many brain pathologies, including hydrocephalus, Alzheimer’s disease, and cognitive dysfunction, are associated with the dysregulation of CSF production or a loss of the blood–CSF barrier integrity [11, 12]. Thus, the ChP has the potential to serve as a gateway for a wide range of hematogenous pathogens.

Numerous pathogens, including viruses, can utilize the blood–CSF barrier as an entry route. Microbes can cross the blood–CSF barrier through two main mechanisms: transcellular and paracellular neuroinvasion. Transcellular neuroinvasion involves the passage of pathogens through the barrier cells themselves (with or without causing infection), with the barrier integrity remaining intact. Conversely, paracellular neuroinvasion involves the translocation of pathogens through the space between the barrier cells. This transport is often facilitated by inflammation or infection–induced cell death, which results in barrier breakdown and allows passive pathogen diffusion into the CNS. A subtype of paracellular neuroinvasion, known as the Trojan horse mechanism, involves barrier crossing by pathogens that first infect the CNS–infiltrating immune cells. Studies using *in vivo* models have shown that human polyomavirus (JCPyV) and human immunodeficiency virus–1 (HIV-1) employ the Trojan horse strategy [13, 14], whereas human herpes simplex virus type 1 (HSV-1) and severe acute respiratory syndrome coronavirus 2 (SARS-CoV-2) damage the epithelial barrier, while the Chikungunya virus enters without disturbing it [15, 16, 17]. For orthoflaviviruses, previous studies have demonstrated ChP susceptibility involving WNV infection in wildtype mice and ZIKV and Langat virus infections in severely immunocompromised mice, indicating the potential for the ChP to serve as a pathogen gateway [18, 19, 20]. However, many of these medically important orthoflaviviruses do not have rodents as their natural hosts, and the mechanism by which they enter the CNS in these studies remains unclear. Therefore, there is a need to understand the role of ChP in the susceptibility, responses, and mechanisms underlying orthoflavivirus neuroinvasion in human models.

Recent developments in ChP organoid models derived from human pluripotent stem cells have led to the creation of microphysiological systems that recapitulate the critical physiology and functions of native ChP [16, 21]. These models exhibit transcriptome and secretome profiles similar to those of human ChP, and their epithelial cells can form the tight junctions essential for maintaining barrier integrity. Notably, the epithelial cells secrete a CSF–like fluid into a self–contained cyst within the organoid. Consequently, these organoids represent a critical platform for *in vitro* investigation of human viral infection and pathogenesis at the blood–CSF barrier, overcoming the inherent interspecies differences of animal models and ethical limitations of human models.

In the present study, we established ChP organoids (ChPOs) derived from inducible pluripotent stem cells (iPSCs) to model the human blood–CSF barrier and validate their functional characteristics. We found that upon infection with various neurotropic orthoflaviviruses, the ChPOs exhibited differential susceptibility, with the greatest susceptibility found for the pathogenic JEV. The organoids enabled an investigation into the intrinsic innate immune response triggered by virus infection. They demonstrated a robust interferon (IFN) response following infection, but they secreted JEV particles into the CSF–like fluid without compromising the epithelial barrier function. This intriguing finding suggests a possible transcellular mechanism for JEV neuroinvasion via the blood–CSF barrier.

## Materials and Methods

### iPSC culture

The human iPSC experiment was approved by the Ethical Committee, Faculty of Ramathibodi Hospital, permit number MURA2022/321. The iPSCs were derived from CD34^+^ hematopoietic stem cells from a healthy Thai donor (clone MURAi003-A, [22]) and from ATCC-BYS0113 (#ACS-1027, ATCC). The iPSCs were cultured on Corning® Matrigel® Growth Factor Reduced (GFR) Basement Membrane Matrix–coated plates at 10 µg/cm^2^ (# 354230, Corning). The cells were dispensed in mTeSR™ Plus medium (#1000274, Stem Cell Technologies) supplemented with 10 µM Y-27632 ROCK Inhibitor (#72302, Sigma–Aldrich) for 24–48 h post–seeding. The culture medium was then replaced with fresh mTeSR medium without a Rock inhibitor every other day. The iPSCs were routinely passaged every 4–6 days by washing with Dulbecco’s phosphate–buffered saline without calcium and magnesium (DPBS, #37350, Stem Cell Technologies) and dissociating with 0.5 mM EDTA (#E7174-1-0500, QReC). Cells were maintained under hypoxic conditions (5% CO_2_ and 3% O_2_) at 37°C.

### ChPO differentiation and maturation

The iPSCs were differentiated into ChPOs using the STEMdiff™ Choroid Plexus Organoid Differentiation Kit (#100-08240, Stem Cell Technologies) and STEMdiff™ Choroid Plexus Organoid Maturation Kit (#100-0825, Stem Cell Technologies) following the protocol stated in Chew et al. [23]. The organoids were maintained in an ultra–low binding culture plate with orbital shaking at 100 rpm and cultured under normoxic conditions (5% CO_2_ in atmospheric O_2_ concentration) at 37°C. The organoids were allowed to mature for 80 days before being used in the infection experiments.

### Viruses and cell lines

JEV strain Nakayama (received from US–AFRIM), JEV strain SA14-14-2 (obtained by virus culture in Vero cells from CD.JEVAX), DENV2 strain US/BID-V594/2006 (# NR-43280, BEI resources), and Zika virus strain PRVABC59 (#NR-50240, BEI resources) were used for the infection experiments. Viruses were amplified in an *Aedes albopictus* (C6/36) cell line (ATCC CRL-1660) in Leibovitz’s L15 medium (#L4386, Sigma–Aldrich) supplemented with 2% fetal bovine serum (#A5256701, Gibco), 2% tryptose phosphate broth (#T9157, Sigma–Aldrich), 1× L–glutamine (#SH30034.01, HyClone), and 1× Pen/Strep (100 U/mL of penicillin and 100 µg/mL of streptomycin, #SV30010 and HyClone) and maintained at 5% CO_2_, 28°C. The JEV–infected supernatant was harvested on day 4 post–infection (pi), whereas the DENV2– and ZIKV–infected supernatants were harvested on days 7–9 pi. All infectious work was performed in a certified biosafety–level 2 lab with approval from the Ramathibodi Hospital Institutional Biosafety Committee (RAMA-IBC 2023-001).

### Plaque assays

Infectious virus titers in the supernatant were determined using plaque assays. Briefly, the virus supernatant was diluted 10–fold in Minimum Essential Medium (MEM, #41500-043, Gibco) without fetal bovine serum (FBS) supplementation and inoculated onto BHK–21 cells (#ATCC-CCL10, ATCC). The cells were allowed to absorb the infectious viruses for 1 h at 5% CO_2_, 37°C and. After absorption, the cells were overlaid with MEM supplemented with 2% FBS and 1% methylcellulose (#M0512, Sigma–Aldrich) and incubated at 5% CO_2_, 37°C for 3, 5, and 7 days for JEV, ZIKV, and DENV2, respectively. The cells were then stained with 0.1% crystal violet (#61135, Sigma–Aldrich) in 20% ethanol (#B6531–03, Fulltime) overnight at room temperature. The stained cells were then washed with water until clear plaques appeared in the positive control wells. The plaques were counted, and the infectious virus titers were calculated as plaque forming units per mL(PFU/mL).

### ChPO infection experiments

The ChPOs were infected with JEV, ZIKV, and DENV orthoflaviviruses. Viruses were added to 1 mL of the maturation medium from the STEMdiff™ Choroid Plexus Organoid Maturation Kit at the following multiplicities of infection (MOIs): 10^3^ or 10^5^ PFU for JEV strain Nakayama and 10^5^ PFU for JEV strain SA–14-14-2, ZIKA, and DENV2. The ChPOs were allowed to absorb the virus supernatant for 4 h, and then the medium was replaced with 1 mL fresh maturation medium. During the course of infection, the virus supernatant was collected daily for plaque assays, and the cultures were supplemented with fresh medium. The infected organoids were cultured for up to 96 h post infection (hpi).

### RNA extraction and RT–qPCR

The ChPOs were washed twice with DPBS then collected in RNAlater Solution (#AM7021, Invitrogen) and stored at -80°C for future RNA extraction. For RNA extraction, the organoids were transferred into the RNA lysis buffer supplied with the Quick–RNA MiniPrep kit (#R1054, Zymo Research) and RNA was extracted following the manufacturer’s protocol. The RNA was eluted from the kit columns by adding 50 µL of UltraPure Distilled Water nuclease–free (#10977-015, Invitrogen). The eluted RNA was stored at -80°C for further use.

To perform RT–qPCR, first–strand cDNA synthesis with random hexamers was performed using SuperScript III First–Strand synthesis system for RT–PCR (#18080-051, Invitrogen). Cellular gene expression was quantified using LUNA Universal qPCR Master Mix (#M3003L, New England Biolabs) according to the manufacturer’s protocol. The primers used in the experiments are shown in Supplementary file 1, Table S1. The experiments were run on a CFX96 Real–Time System (Biorad). Relative mRNA expression levels were normalized to GAPDH using the ΔΔCt method [24]. The results were visualized using GraphPad Prism V.6 (GraphPad).

### Tissue sectioning, staining, and imaging

The ChPOs were fixed with 4% paraformaldehyde (#1003550033, Sigma–Aldrich) at room temperature for 15 min, washed 3 times in DPBS, and then dehydrated in 30% w/v sucrose solution at 4°C. The tissue was snap–frozen in Tissue–Tek O.C.T. compound (#4583, Sakura) using liquid nitrogen and then sectioned at a thickness of 20 µm using a rotary microtome cryostat.

The immunofluorescence assay was performed by permeabilizing the tissue sections and blocking them in 10% v/v goat serum and 0.2% v/v TritonX-100 in DPBS for 1 h at room temperature. The tissues were then immunolabeled with primary antibodies at the following dilutions: anti–TTR 1:200 (#703278, Invitrogen), anti–Zo1 1:500 (#AB96587, Abcam), GFAP 1:500 (#AB190288, Abcam), TUBB 1:100 (#sc-51670, Santa Cruz), and 4G2 (hybridoma supernatant, neat, produced in–house). All antibodies were diluted in 0.5% v/v TritonX-100 in PBS. The incubations were performed at room temperature for 2 h, followed by 3 PBS washes. For staining, the following secondary antibodies were used: anti–rabbit Alexa Fluor 488 (#4412SS, Cell Signaling) or anti–mouse Alexa Fluor 555 (#4409S, Cell Signaling) diluted at 1:500 for 1 h at room temperature in the dark. The sections were then counterstained with DAPI (#10236276001, Sigma–Aldrich) for 5 min and mounted with Antifade Mounting Medium (#H-1200, Vectashield). Confocal fluorescence microscopy was performed using a Zeiss 900 Airyscan 2 confocal microscope (Zeiss) using Zen 3.4 blue ver. 3.4.91.00000, and the images were processed using Image J (NIH) imaging software.

### RNA sequencing and transcriptomics analysis

RNA sequencing was performed with 3 pooled samples of 3 ChPOs per condition, with the following treatments: JEV strain Nakayama infected organoids (10^5^ PFU for 24 h), Poly(I:C) (#tlrl-pic, InvivoGen) treated organoids (20 µg/mL for 24 h), and mock–treated organoids. Total RNA was extracted from the organoids as mentioned above using the Quick–RNA Miniprep Kit (#R1054, Zymo Research) in accordance with the manufacturer’s instructions. RNA integrity was assessed using a 4200 TapeStation (Agilent Technologies, Palo Alto, CA, USA) and a 1% agarose gel. A 300 ng sample of total RNA with an RIN value above 6 was used to construct cDNA libraries using the polyA mRNA isolation kit (VAHTS Universal V8 RNA–seq Library Prep Kit). Sequencing, which was performed on the Illumina NovaSeq 6000 instrument according to manufacturer’s instructions (Illumina, San Diego, CA, USA), generated paired–end 150 bp reads (PE150, 2 × 150 bp).

The sequencing reads were mapped to the human reference genome (hg38) using HISAT2 [25], and gene–level read counts were generated with Rsubread [26]. Differential expression analysis was carried out in R (v4.4.1) using the DESeq2 package [27] based on triplicate biological samples. Genes were considered differentially expressed genes (DEGs) when they exhibited a log2–fold change greater than 1 or less than −1 (|log_2_ fold change| ≥ 1) with an adjusted *p*-value < 0.05. Gene Set Enrichment Analysis (GSEA) was performed with the hallmark gene sets from MSigDB [28]. Genes were ranked in descending order and analyzed with the clusterProfiler R package [29]. Pathways were considered significantly enriched when the Benjamini–Hochberg adjusted *p*-value was below 0.05.

### Treatment with interferon receptor antibodies

To investigate the role of type–I and type–III IFN responses to JEV infection, ChPOs were treated with 1 µg/mL anti–hIFNAR1 (Anifrolumab) antibody (#hifnar–mab1, InvivoGen) and/or 5 µg/mL of an anti–hIFNLR3 (Anti-hIL-28b) neutralizing mAb (#mabg-hil28b-3, InvivoGen) 24 h prior to JEV infection. Antibodies were supplemented in the culture medium throughout the experiment, both during the virus absorption and during the 3–day incubation period.

### Evaluation of ChPO barrier integrity

To determine the size selectivity of the ChPO barrier, the mature organoids (Day 80) were incubated with 0.5 mg/mL of a cocktail containing fluorescein (332.29 Da; #F1300, Invitrogen), Cascade Blue™ dextran, 3000 MW (#D7132, Invitrogen), and tetramethylrhodamine dextran, 70,000 MW (#D1818, Invitrogen) diluted in DPBS. After a 2 h incubation, the supernatant (medium) was collected, and ChPOs were given five min DPBS washes. The CSF–like fluid was collected using an insulin needle [23]. The fluorescence intensities of the medium and CSF–like fluid were measured at excitation/emission (Ex/Em) 494/521 nm, Ex/Em 400/420 nm, and Ex/Em 555/580 for the 0.332, 3, and 70 KDa dyes, respectively, using Cytation 5 (Agilent BioTek).

### Viability assays

The viability of the ChPOs was assessed using a resazurin assay (#ab129732, Abcam). In brief, the organoids were incubated in the maturation medium mixed with 1× resazurin for 4 h. After incubation, 100 μL of culture medium was transferred to a black 96–well plate, and the fluorescence intensity was measured at Ex./Em. 560/590. ChPOs without treatment were used as viable organoid controls, and ChPOs treated with neat DMSO for 10 min were used as killed controls.

## Results

### Establishment and characterization of ChPOs derived from human iPSCs

We generated ChPOs using CD34^+^–derived iPSCs obtained from a healthy Thai donor at Ramathibodi Hospital (MURAi003-A) (Fig. 1A) [22]. The ChPOs were first induced to form embryoid bodies (EB), followed by differentiation into neuroepithelium in Matrigel. The organoids were then allowed to mature in the maturation medium. The successfully formed ChPOs showed cysts filled with CSF–like fluid by days 25–35, closely mimicking the fluid–secretion function of the ChP epithelium *in vivo*. A study by Pellegrini et al., who established the protocol for generating ChPO used in our study, also demonstrated that the protein profile in the CSF–like fluid from organoids older than 60 days was similar to that of the pediatric to adult stages [21], and a study by Qiao et al. reported that the 80–day–old ChPO epithelium displayed a columnar structure [30]. Consequently, we allowed our ChPOs to mature for 80 days before using them in the experiments. The sizes of ChPOs varied, with the majority of them growing to sizes of up to 2–3 mm in diameter, while about 5% of them were larger than 5 mm in diameter and contained large and harvestable fluid–filled cysts.

**Fig 1.**
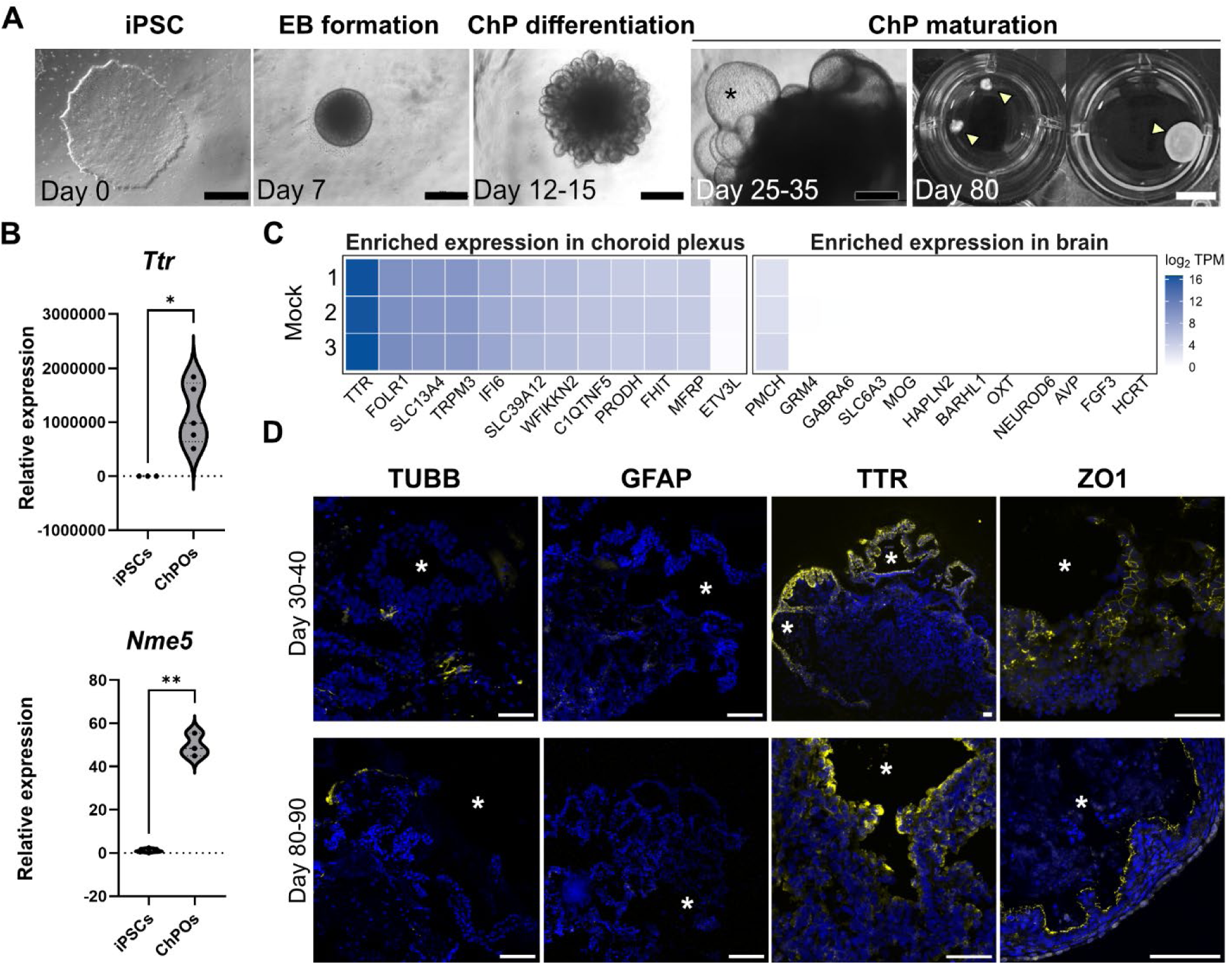
Generation and characterization of iPSC–derived ChPOs. **(A)** Representative images of ChPO differentiation and maturation from MURAi003-A. A generation of ChPO was initiated with EB formation. Subsequent ChP differentiation occurred by the formation and expansion of neuroepithelium. At around days 25-35, the organoids began to form fluid–filled cysts (marked with *) containing CSF–like fluid. The ChPOs were allowed to mature up to 80 days (marked with yellow ▴). Black scale bars = 500 µm and white scale bar = 1 cm. **(B)** Expression of mature ChP marker genes *Ttr* and *Nme5* in iPSCs compared with 80–day–old ChPOs. Mean ± SD; * *p* < 0.05; ** *p* < 0.01; Unpaired t test with Welch’s correction. **(C)** Heatmap of gene expression found in the 80–day–old ChPOs. The gene lists were selected from the highest level of enriched expression in ChP and in the brain from the Human Protein Atlas. The expression level was displayed in Log2 TPM. **(D)** Immunofluorescence staining of 30-40–day–old and 80-90–day–old ChP with markers for neurons (TUBB), astrocytes (GFAP) and ChP epithelium (TTR and ZO1) in yellow and counterstained with DAPI in blue. Asterisks (*) indicate fluid–filled cysts areas. Scale bars = 100 µm.

We confirmed successful ChPO differentiation by quantifying the expression of the mature ChP marker genes, transthyretin (*Ttr*) and nucleoside diphosphate kinase homolog 5 (*Nme5*), and found high expression of these genes in the 80–day–old ChPOs (Fig. 1B) [21]. Transcriptomic profiling also showed high expression of the ChP–specific gene set compared to a brain–specific gene set acquired from the Human Protein Atlas Database [31] (Fig. 1C). Immunofluorescence analysis revealed minimal expression of β–tubulin (TUBB), a marker of mature neurons, and no expression of glial fibrillary acidic protein (GFAP), a glial cell marker (Fig. 1D). In contrast, immunofluorescence staining for TTR and the tight junction protein ZO1, which are both markers of ChP epithelium, was observed at the rims of the fluid–filled cysts. Similar ChP differentiation and marker localization were observed from ChP derived from another CD34^+^–derived iPSC line purchased from ATCC (BYS0113; Supplementary file 1, Fig. S1). Taken together, these results demonstrated the consistent generation of ChPOs that contained epithelial cells at the periphery of the fluid–filled cysts, thereby closely mimicking the morphology of the *in vivo* ChP.

### ChPOs are susceptible to neurotropic orthoflavivirus infection

Several orthoflaviviruses, such as JEV, ZIKV, and DENV, are neuroinvasive and display neurotropism toward cells in the CNS. We explored whether ChPOs could be targeted by and serve as replication sites for neurotropic orthoflaviviruses by infecting ChPOs with these viruses and quantifying the titer of the infectious virus particles in the supernatant (Fig. 2A). The plaque assay results showed that ChPOs were highly susceptible to the pathogenic JEV strain Nakayama. The ChPOs from MURAi003-A produced and released a large amount of infectious virus progeny into the supernatant, plateauing at around 10^5^ PFU/mL by 48 h post–infection (hpi) (Fig. 2A). The ChPOs from BYS0113 gave similar results but were slightly more susceptible to the infection, plateauing at 10^6^ PFU/mL by 72 hpi (Fig. 2A). The ChPOs supported lower replication of JEV strain SA14-14-2, the attenuated vaccine strain that is non–neuroinvasive [32]. Although we infected the virus at 100–fold higher titer than we used with the JEV strain Nakayama, the JEV strain SA14-14-2 reached a maximum titer of only 10^3^ PFU/mL (Fig. 2B). In addition to their susceptibility to JEV, the ChPOs showed moderate susceptibility to ZIKV infection and gave a titer similar to that achieved with JEV strain SA14-14-2 (Fig. 2C). The ChPOs were least susceptible to DENV2 infection (Fig. 2D). One of the three organoids had an abortive infection, while the other two produced infectious particles at levels below 10^3^ PFU/mL. Since the ChPOs were most susceptible to the JEV strain Nakayama, subsequent experiments were performed using this strain, unless otherwise specified.

**Fig 2.**
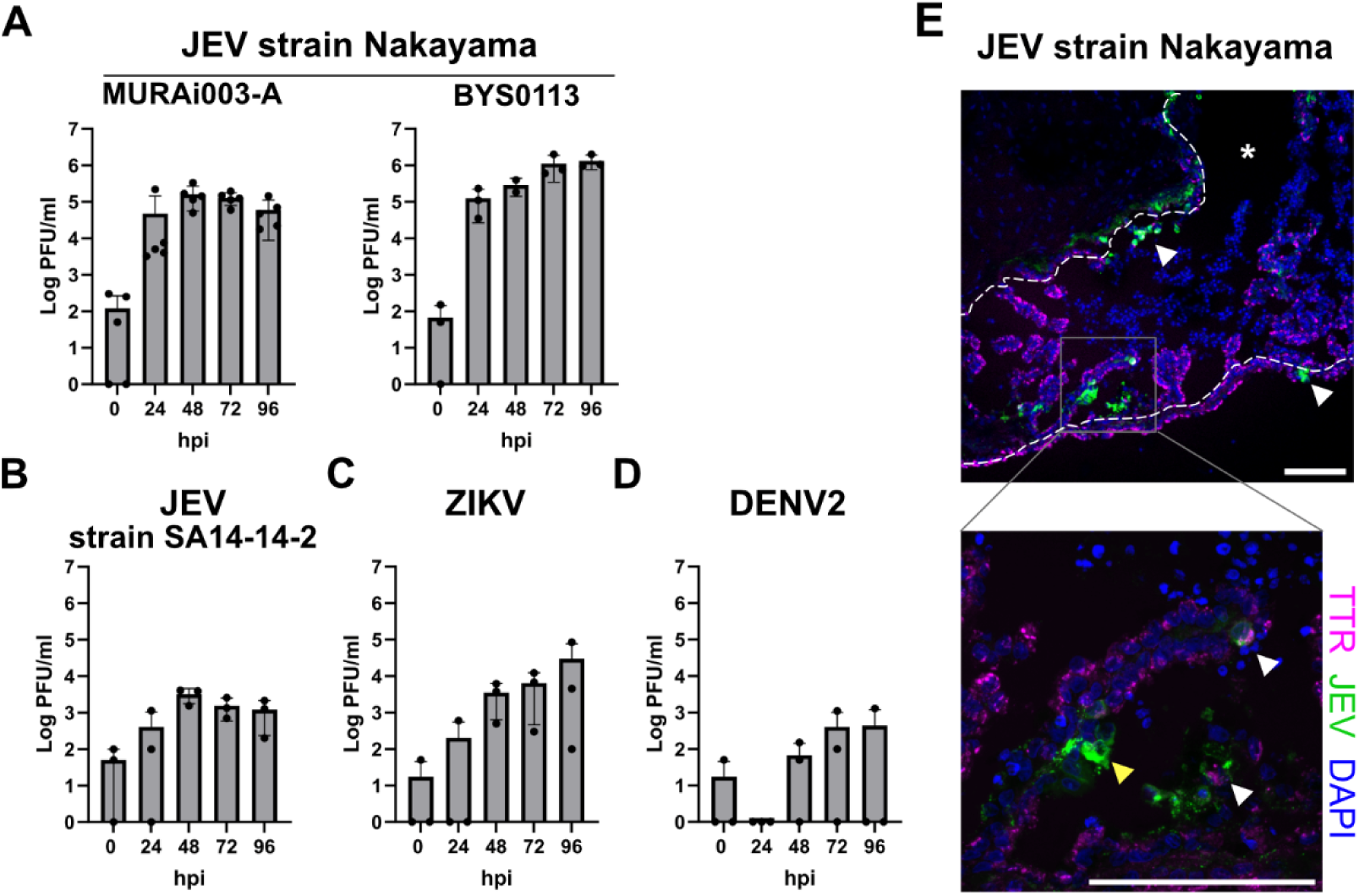
ChPOs susceptibility and tropism to orthoflavivirus infections. **(A-D)** Quantification of virus titers in the culture supernatant by plaque assay. Eighty–day–old ChPOs derived from MURAi003-A were infected with **(A)** JEV strain Nakayama at 10^3^ PFU, **(B)** JEV SA14-14-2 at 10^5^ PFU, **(C)** ZIKV strain PRVABC59 at 10^5^ PFU and **(D)** DENV2 strain US/BID-V594/2006 at 10^5^ PFU. Cultured supernatant was collected for plaque titration and replaced daily up to 96 hpi. The titer on hour 0 indicates the residual amount of the input virus in the supernatant. Increased titers at the following timepoints indicate active virus infection. **(E)** Representative image showing co–staining of JEV strain Nakayama envelop protein with 4G2 antibody (green) and ChP epithelial cells marker TTR (magenta) in JEV–infected ChPO. Eighty–day–old ChPO was infected with JEV strain Nakayama at 10^5^ PFU and the organoid was collected for fixation and cryopreservation at 96 hpi. The organoid was sectioned at a thickness of 20 μm, stained with antibodies, and counterstained with DAPI (blue). White arrowheads indicate cells that co–stained with 4G2 and TTR, indicating infected ChP epithelial cells, while yellow arrowheads indicate infected cells that are not epithelial cells. Broken lines show the border of the fluid–filled cyst and the luminal side is marked with *. The boxed area is a magnified image from the top panel. Scale bar =100 μm.

We then investigated the cellular tropism of JEV infection within the ChPO by immunofluorescence staining with 4G2, an antibody against pan–flaviviral envelope protein, together with anti–TTR, a ChP epithelial marker (Fig. 2E). We detected JEV infection in cells both with and without TTR expression, suggesting that JEV has tropism toward both ChP epithelial cells and other cell types within the ChPOs.

### ChPOs display a functional antiviral response against virus infection

The ChP acts as a barrier that protects the brain from harmful substances, including viruses, circulating in the periphery. Apart from serving as a physical wall, ChP cells may also possess intrinsic biological mechanisms that enable them to sense and counteract virus infections. We investigated how iPSC–derived ChPO would respond to invading neurotropic viruses, particularly JEV, using bulk RNA sequencing and transcriptomics analysis. We examined the transcriptomic profiles of ChPOs after treatment with Poly(I:C), a synthetic double–stranded RNA analog that mimics viral molecular patterns, or after infection with JEV, and compared the profiles to those of uninfected (mock) ChPO controls (Fig. 3A). Principal component analysis (PCA) of the transcriptome revealed a clear separation of samples from the different treatment groups (Fig. 3B). The transcriptomic profiles of Poly(I:C)–treated ChPOs showed a greater divergence from the mock controls than did the profiles of the JEV–infected ChPOs. This finding indicates that a more robust transcriptional changes was elicited in ChPOs by Poly(I:C) treatment than by JEV infection in this system.

**Fig 3.**
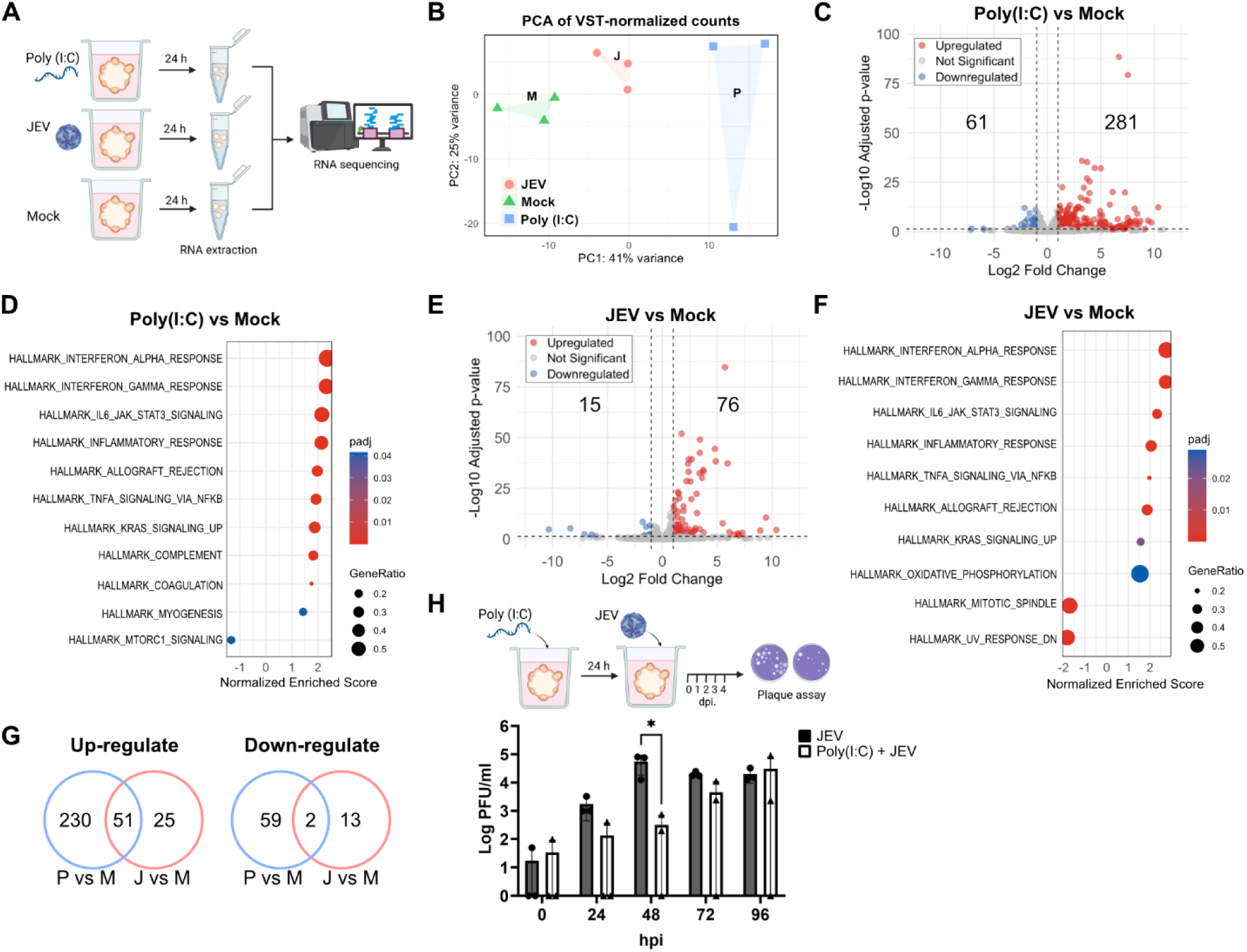
Transcriptomic analysis of Poly(I:C)–treated and JEV–infected ChPOs compared with the mock ChPOs. **(A)** A schematic shows the experimental design. The ChPOs were treated with 20 µg/mL Poly(I:C) or infected with 10^5^ PFU of JEV strain Nakayama or mock–treated for 24 h. Three organoids were pooled for each sample, and three samples per condition. Samples were subjected to bulk RNA sequencing and transcriptomic analysis. **(B)** Principal Component Analysis (PCA) of transcriptomes of Poly(I:C)–treated, JEV–infected, and mock CHPOs. **(C)** A volcano plot comparing DEGs of Poly(I:C) treated vs. mock ChPOs. **(D)** GSEA of DEGs of Poly(I:C) treated vs. mock ChPOs. The dot plot shows the top enriched hallmark gene sets identified by GSEA. The dot size is proportional to the gene ratio, and the color indicates adjusted *p*-value. **(E)** A volcano plot comparing DEGs of JEV–infected vs. mock ChPOs. **(F)** GSEA of DEGs of JEV–infected vs. mock ChPOs. **(G)** Venn diagrams showing the overlap of upregulated or downregulated DEGs in JEV–to–mock (J vs. M) comparisons to Poly(I:C)–to–mock (P vs. M) comparisons. **(H)** A schematic illustrates the experimental design and a plaque titration result of the supernatant collected from ChPOs infected with 10^5^ PFU JEV strain Nakayama, with and without 20 µg/mL Poly(I:C) pretreatment. Noted that ChPO supernatant was supplemented with Poly(I:C) only during the pretreatment but not at time of infection and post–infection. Mean ± SD; * *p* < 0.05; Two–way ANOVA with Sidak’s multiple comparisons test.

The transcriptomic profiles of Poly(I:C)–treated vs. Mock ChPOs revealed 342 differentially expressed genes (DEGs; 281 upregulated and 61 downregulated; |log_2_ fold change|≥ 1 with adjusted *p*-value < 0.05) (Fig. 3C, Supplementary file 2). Gene Set Enrichment Analysis (GSEA) revealed that the top four enriched Hallmark gene sets, all involved in immune responses, were interferon_alpha_response, interferon_gamma_response, IL6_Jak_Stat3 signaling, and inflammatory_response gene sets (Fig. 3D).

Similarly, the transcriptomic profiles of JEV–infected vs. mock ChPOs showed 76 upregulated and 15 downregulated DEGs (Fig. 3E, Supplementary file 3). GSEA revealed the same top four pathways as observed in Poly(I:C)–treated ChPO samples (Fig. 3F). We found that genes that were commonly upregulated in both Poly(I:C)–treated and JEV–infected ChPOs were genes involved in interferon (IFN) induction through the RIG–I–like receptor pathway (*Ifih1* and *DDXH58*), IFN response pathway (*Stat1* and *Irf7*), and interferon–stimulating genes (ISGs; such as *Oas1*, *Mx1*, and *Ifit3* etc.) (Fig. 3G). Conversely, two commonly downregulated genes were LOC101927314 and *Nup210l*. These genes have no known role in responses to virus infection, but *Nup210*, the paralog of *Nup210l*, is known to play a role in inhibiting HIV–1 infection [33]. These results demonstrate that ChPOs are able to elicit interferon responses upon JEV infection.

We tested whether the antiviral response stimulated by Poly(I:C) was functional and able to prevent future virus infection by pretreating ChPOs with Poly(I:C) for 24 h, followed by infection with JEV. The plaque titration results indicated slower infection kinetics for the JEV infection with prior Poly(I:C) treatment, and significantly lower virus amounts appeared at 48 h post–infection (hpi) in the supernatant of ChPOs pretreated with Poly(I:C) than without pre–treatment (Fig. 3H). In fact, one of the three pretreated organoids escaped JEV infection throughout the experiment. However, we did not supply fresh Poly(I:C) in the media after the time of infection. Therefore, the inhibitory effect of the Poly(I:C) to suppress virus infection subsided on the later days and the virus was able to replicate at high levels by 96 hpi. Taken together, the transcriptomic and infection results suggest that the ChPOs are equipped with functional antiviral machinery that is induced by the IFN pathway.

### Type–I IFN responses play a dominant role in inhibiting JEV infection

We investigated the IFN responses of ChPOs to JEV infection further by first reconstructing the IFN pathways by overlaying the transcripts from our transcriptomic datasets onto the IFN induction and IFN response pathways (Fig. 4A, indicated in black font). In the figure, the genes that were differentially expressed during JEV infection are indicated in bold font, with upregulation or downregulation indicated by upward or downward triangles, respectively (Fig. 4A, bold). Interestingly, the upregulated DEGs can be mapped to key proteins in the IFN induction pathway through the RIG–I–like receptor (RLR) and the IFN response through the JAK–STAT pathway. Ultimately, these signaling cascades lead to the increased expression of several chemokines and ISGs, which possess immune cell–attractant and antiviral properties, respectively.

**Fig 4.**
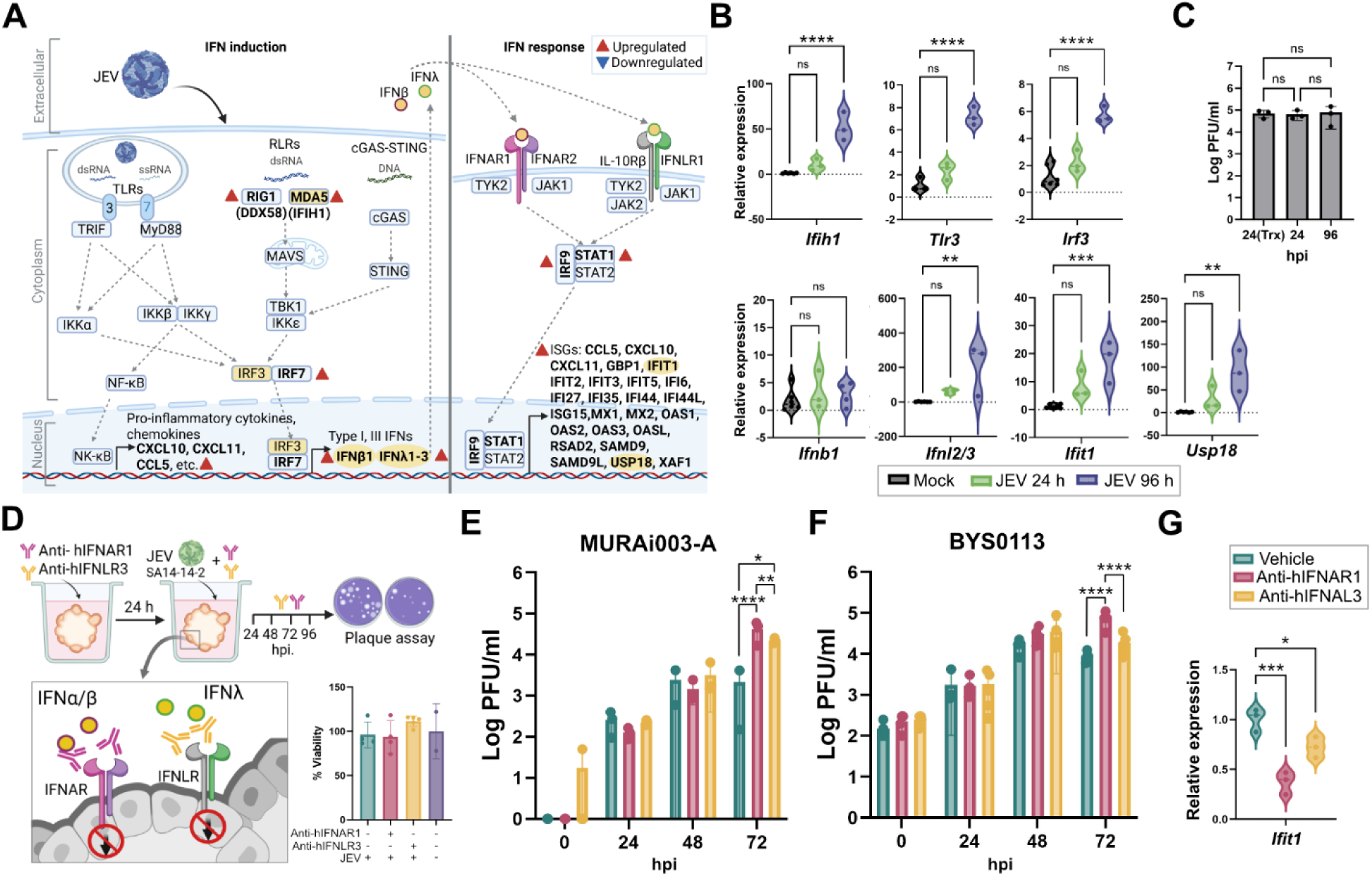
IFNs response in JEV–infected ChPOs. **(A)** A schematic illustrates the canonical protein components of the IFN induction and response pathways, overlaid with data from our bulk transcriptomic analysis. Gene transcripts corresponding to the pathway proteins are categorized by font: “black” for detected transcripts with no significant differential expression; “blue” for transcripts that were not detectable; and “bold” for DEGs. Significant upregulation of DEGs is specifically marked by red arrowheads (▴). Components highlighted in yellow indicate the selected genes used for independent RT–qPCR validation, as detailed in Fig. B. **(B)** Relative expression of the selected genes from the independent ChPO infection experiments quantified by RT–qPCR. The ChPOs were infected with 10^5^ or 10^3^ PFU of JEV strain Nakayama and the supernatants were collected at 24 and 96 hpi, respectively. **(C)** Plaque assay results comparing JEV titers from the supernatant of JEV–infected ChPOs from the transcriptomics experiment (24 Trx) harvested at 24 hpi, to the titers from the RT–qPCR experiment harvested at 24 and 96 hpi. **(D)** A schematic illustrates the experimental design and viability assay of the anti–IFN receptor antibody treatment and JEV infection. ChPOs were pre–treated with 1 µg/mL anti–IFNAR1 or 5 µg/mL anti–IFNLR3 at 24 h prior to infection with 10^5^ PFU of JEV strain SA14-14-2. Supernatants were collected daily up to 72 hpi for plaque titration. Fresh media supplemented with the antibodies were replaced once daily. The viability assay was performed on the ChPOs infected for 72 h (treated with antibodies for 96 h). The result was reported as a percentage relative to the mock. **(E-F)** Plaque titration results of the supernatant collected from ChPOs derived from MURAi003-A (E) or BYS0113 (F), treated with anti–IFN receptor antibodies or vehicle and infected with JEV as demonstrated in Fig. D. **(G)** Relative expression of *Ifit1* quantified by RT–qPCR. Total RNA was extracted from the ChPOs (BYS0113) tissue infected with JEV strain SA14-14-2 for 72 hpi (treated with antibodies for 96 h). Figures E-G share the same graph legend as displayed in G. Statistics: (B, D and G) Mean ± SD; One–way ANOVA with Dunnett’s multiple comparisons test. (C) Mean ± SD; One–way ANOVA with Tukey’s multiple comparisons test. (E and F) Mean ± SD; Two–way ANOVA with Tukey’s multiple comparisons test. * *p* < 0.05; ** *p* < 0.01, *** *p* < 0.005, **** *p* < 0.001.

We then validated the ChPO responses to JEV infection demonstrated in the transcriptomic data by performing RT–qPCR on RNA samples harvested from independent experiments (Fig. 4B). We also performed this evaluation on samples infected with JEV at 10^5^ and 10^3^ PFU and harvested them at 24 and 96 hpi, respectively, to capture the expression of both early and late responses. Notably, these samples had comparable virus titers in the supernatant at both timepoints when compared with the samples used for transcriptomic analysis (Fig. 4C). Seven genes belonging to various steps in IFN induction (*Ifih1, Trl3*, *Irf3*, *Ifnb1,* and *Ifnl2/3*) and IFN responses (*Ifit1* and *Usp18*) were selected for further validation (Fig. 4A, highlighted in yellow, and Fig. 4B). The RT–qPCR results showed that most of these genes, except for *Ifnb*1, showed a similar trend of responses when compared to the transcriptomic data, which increased in expression upon JEV infection. Although the expression levels of these genes were not significantly upregulated at 24 hpi, they became significantly upregulated by 96 hpi. The *Ifnl2/3* gene (encoding IFNλ, the type–III IFN that predominantly responds in epithelial cells) exhibited the highest fold change, increasing over 150–fold compared to the mock treatment (Fig. 4B). Conversely, the *Ifnb1* gene, which encodes for the canonical antiviral type–I IFN, IFNβ, remained unchanged throughout the course of the infection.

The robust upregulation of IFNλ in JEV–infected ChPOs, along with the tissue–specific activity of type–III IFNs in epithelial barriers, led us to hypothesize that type–III IFNs may play a more critical role than type–I IFNs in this infection model. We tested this hypothesis by validating the function of both IFNs by treating the ChPOs with antibodies against type–I and type–III IFN receptors (anti–hIFNAR1 and anti–IFNLR3, respectively) (Fig. 4D). The antibodies were added to the organoid supernatant 24 h prior to infection and throughout the course of infection. In this functional assay, we infected ChPOs with the JEV strain SA-14-14-2, a strain that normally achieves a moderate infection titer under IFN–competent conditions. This ensured that the assay operated within a measurable increase in viral replication upon treatment with anti–IFN antibodies, thereby avoiding a ceiling effect that would have masked the true protective efficacy of the IFN pathway. We anticipated that receptor inhibition would impair the IFN response, resulting in an increased viral titer, especially for type–III IFN. The plaque titration data revealed that inhibition of IFN responses with IFN receptor antibodies resulted in increases of JEV infectivity than was achieved using vehicle treatment. Furthermore, the inhibition of type–I IFN response resulted in a significantly higher JEV titer than was observed following type–III IFN inhibition (Fig. 4E and 4F). Inhibition of IFNs also resulted in a reduction in *Ifit1* expression, an antiviral ISG, confirming the impaired antiviral state and a correlation with increased infectivity.

Taken together, the results demonstrated that JEV infection can induce an IFN response in the ChPOs. This activity plays an important role in controlling JEV infection, with type–I IFN appearing to act as the dominant pathway, providing superior antiviral protection against JEV infection in the ChPOs.

### JEV translocates into the CSF–like fluid without compromising ChPO barrier integrity

Our finding that ChPOs, and especially ChPO epithelial cells, were susceptible to JEV infection (Fig. 2A and 2E) led us to investigate whether the ChP was the potential site of JEV entry into the CSF and subsequently into the CNS. We harvested sufficient CSF–like fluid by utilizing the 80–day–old ChPOs that formed large cysts (accounting for approximately 5% of cultured ChPOs), as this enabled precise CSF fluid collection using an insulin needle (Fig. 5A). Comparison of the viral titer in the supernatant to the titer in the CSF–like fluid inside the organoid lumen revealed comparable viral titers in both fluids (Fig. 5B).

**Fig 5.**
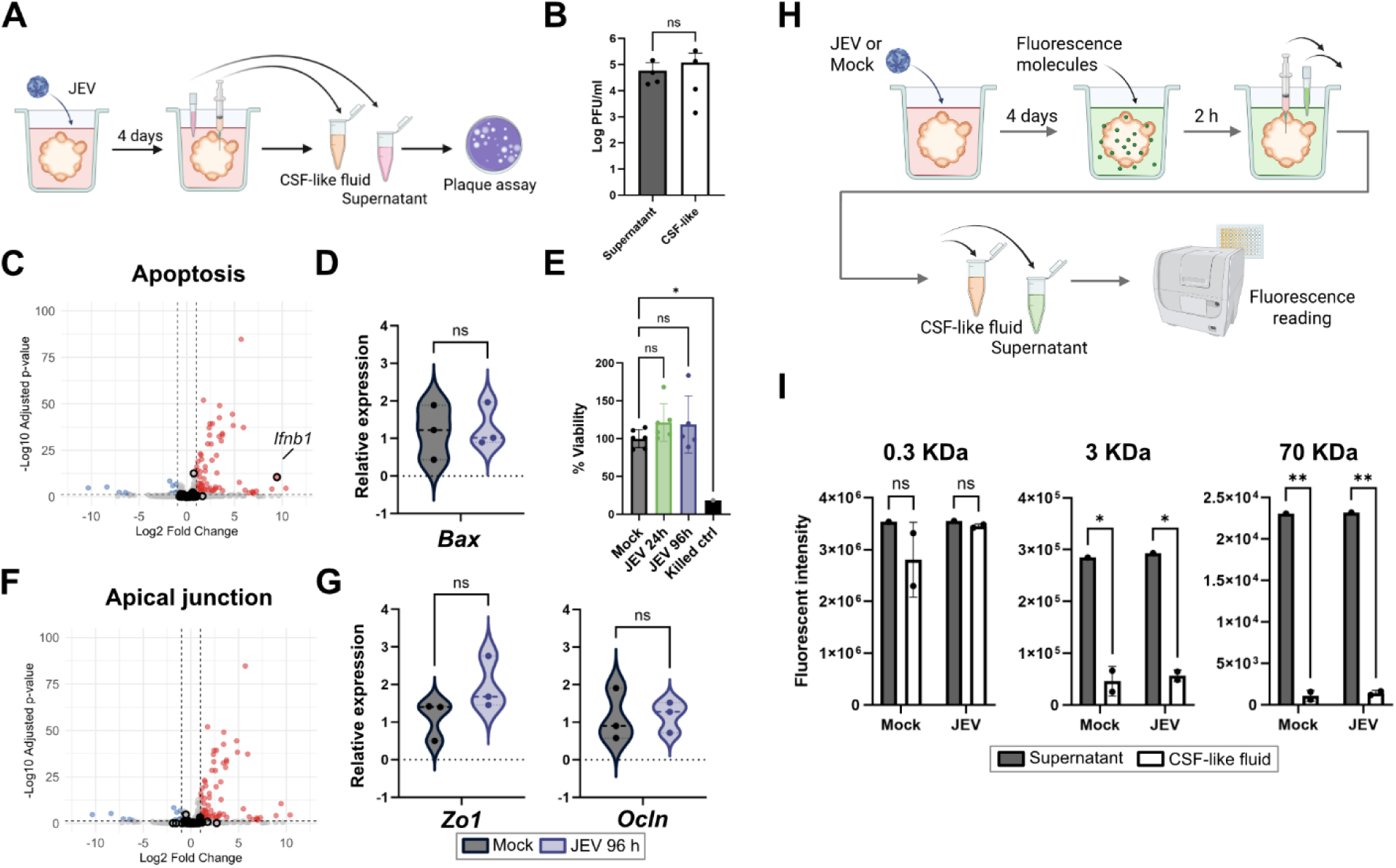
JEV translocates into the CSF–like fluid without causing cell death or barrier leakage. **(A)** A schematic illustrates the experimental design. Eighty–day–old ChPOs with enlarged fluid–filled cysts were infected 10^3^ PFU JEV strain Nakayama. Infectious supernatant and CSF–like fluid were collected and subjected to plaque titration. **(B)** A result of plaque titration of supernatant and CSF–like fluid harvested at 96 hpi. **(C)** A volcano plot comparing DEGs of JEV–infected vs. mock ChPOs. Open circles marking gene members of the Hallmark_Apoptosis pathway. **(D)** Relative expression of *Bax* from the independent ChPO infection experiments quantified by RT–qPCR. **(E)** The viability assay was performed on ChPOs infected with JEV for 24 and 96 hpi, compared to mock (untreated) and killed control. The result was reported as a percentage relative to the mock. **(F)** A volcano plot comparing DEGs of JEV–infected vs. mock ChPOs. Open circles marking gene members of the Hallmark_Apical_Junction pathway. **(G)** Relative expression of *Zo1* and *Ocln* from the independent ChPO infection experiments quantified by RT–qPCR. Figures D and G share the graph legend as displayed in G. **(H)** A schematic illustrates the experimental design for the fluorescent dye passage. At 96 h post–JEV strain Nakayama infection, 0.3, 3, and 70 kDa dye cocktails were added to the infected ChPOs and incubated for 2 h. Supernatant and CSF–like fluid were harvested and measured for the presence of the fluorescent dyes using the Cytation 5 fluorescent reader. **(I)** Fluorescent intensity of 0.3, 3 and 70 KDa dyes measured from the supernatant and CSF–like fluid. Statistics: (B) Mean ± SD; Unpaired t–test with Welch’s correction. (D, E, G) Mean ± SD; One–way ANOVA with Dunnett’s multiple comparisons test. (I) Mean ± SD; Two–way ANOVA with Sidak’s multiple comparisons test. * *p* < 0.05, ** *p* < 0.01.

We then investigated how the virus translocates into the CSF–like fluid, assuming that one possible mechanism of entry could be the paracellular diffusion of virus particles into the organoid through a compromised barrier. Previous studies have shown that DENV and JEV infections lead to apoptotic cell death in the brain endothelial cells and neurons, respectively [34]. Therefore, we first exploited our ChPO transcriptomic profiles to investigate whether JEV infection induced cell death that would compromise the ChPO barrier. Our GSEA analysis of the JEV–infected ChPO transcriptome revealed no significant dysregulation of the hallmark apoptosis pathway (Fig. 3F and 5C). None of the genes within the Hallmark_Apoptosis gene set, except for *Ifnb1*, were differentially expressed following JEV infection. The RT–qPCR result of *Bax*, a key pro–apoptotic factor, further confirmed that apoptotic pathway was not induced in the JEV–infected ChPOs (Fig. 5D). Similarly, none of the genes that encode for the key regulators or key effectors of pyroptosis (*Nlrp3*, *Casp1*, *Trem2*, *Tnf*, *Mmp9*, and Il1b) or necroptosis (Rip1 and Rip3) were differentially expressed, except for *Mlkl*, which encodes for an executioner protein of necroptosis (Supplementary file 3). However, the upregulation of this gene did not contribute to the overall viability of the organoids, as determined by resazurin viability assays conducted both at the early (24 hpi) and late (96 hpi) timepoints (Fig. 5E).

Although the cells at the barrier maintained their viability, the barrier integrity could become compromised if its tight junctions were disturbed. However, our GSEA analysis of the JEV–infected ChPO transcriptome revealed no significant dysregulation of the Hallmark_Apical_Junction, and none of the genes within this Hallmark gene set were differentially expressed (Fig. 3F and 5F). Further RT–qPCR results for genes encoding the tight junction protein Occludin (*Ocln*) and the tight junction–associated protein ZO1 (*Zo1*) also confirmed no dysregulation in the JEV–infected ChPOs (Fig. 5G). Finally, we assessed ChPO barrier integrity by monitoring the trans–epithelial passage of various molecular weights fluorescent dyes (0.321 KDa fluorescein and 3 KDa and 70 KDa fluorescence dextrans; Fig 5H). The results revealed that the cell junctions of both the mock–infected and JEV–infected ChPOs were permissive only to the smallest 0.3 KDa dye but restricted the passage of the 3 KDa and 70 KDa dyes (Fig. 5I). This consistent exclusion profile demonstrated that although JEV can infect ChPO epithelial cells, the infection does not compromise the functional ChP barrier integrity. These findings suggest that JEV entry into the CSF–like fluid does not occur due to barrier leakage, but is instead most likely mediated by the release of viral particles from infected ChP epithelial cells into the CSF–like fluid in the lumen of the ChPOs.

## Discussion

The brain barriers constitute critical anatomical and physiological defense structures, as they serve as the final physical interface restricting virus entry into the CNS and preventing subsequent neurological damage. Several medically important orthoflaviviruses, including WNV, ZIKV, DENV, TBEV, and JEV, are known to be neuroinvasive. Given their systemic hematogenous dissemination and evidence of damage to the BBB, many studies have concentrated on neuroinvasion mechanisms at this site [35]. However, several studies have detected the presence of viruses in the brain parenchyma prior to or concurrent with any observed BBB damage or increased permeability [8, 36, 37, 38, 39]. This evidence suggests that BBB leakage may be a consequence of, rather than the initial cause or site of, neuroinvasion. This possibility underscores the necessity of investigating alternative JEV entry mechanisms.

The blood–CSF barrier has largely been overlooked and underinvestigated. Nevertheless, a growing body of evidence suggests that this barrier has the potential to serve as a site of entry for several viruses, including orthoflaviviruses. For instance, experimental mouse model studies have shown that both ZIKV and WNV were detectable in the ChP following footpad inoculation of *Ifnar^-/-^* and wildtype mice, respectively [18, 20]. Furthermore, WNV was detected in the ChP during a postmortem investigation of an infected mule [39]. In the present study, we modeled this interface in humans by establishing human–iPSC–derived ChPOs, and our results demonstrate that these organoids exhibit varying degrees of susceptibility to JEV, ZIKV, and DENV2 infection.

The differing degrees of ChPO susceptibility may reflect the availability of viral–specific receptors/host factors in ChPO cells. Known host factors for JEV, such as vimentin, glycosaminoglycan (GAGs), glucose–regulated protein 78 (GRP78), and heat shock protein 70 (HSP70), are highly expressed in brain tissues [40, 41, 42]. However, the latter three proteins are also reported to be important host factors for ZIKV and DENV, which were unable to infect well in our system [43, 44]. Therefore, other as yet unknown factors must contribute to susceptibility and require further in–depth studies.

The degree of ChPO infectivity also seems to correlate with the neuroinvasive character of each virus. Consistent with its high clinical neuroinvasion incidence (∼1 in 250) and fatality rate of up to 25% [3], the pathogenic JEV strain Nakayama exhibited superior replication efficiency in ChPOs even when inoculated at 100–fold lower titers than used for the other viruses. Conversely, the attenuated vaccine strain, JEV strain SA14-14-2, which has a documented loss of neurovirulence [45], produced only a moderate infection at 10^5^ PFU. Similarly, ZIKV, primarily known to cause a severe congenital syndrome but milder adult disease [46], also resulted in only moderate infection in our ChPOs at 10^5^ PFU. This low epithelial infectivity aligns with prior findings by Kim et al. [20] that ZIKV has tropism toward pericytes of the ChP (which do not exist in our ChPOs) and traverses through human ChP epithelial cells (HIBCPP) by breaking this barrier but without infecting the ChP epithelial cells. Lastly, DENV2 was the least susceptible virus in our model, with one sample showing no infection at any time point. Neurological manifestations of DENV have been reported in 0.5–46.3% of symptomatic patients [47]. However, this included indirect encephalopathy resulting from other systemic complications, such as metabolic disturbances and hemorrhage, or overdiagnosis due to reliance solely on CSF serology, which could have cross–reactivity due to previous flavivirus exposures. The true neuroinvasion and infection events in DENV are thought to be much lower than the estimated values [48]. Overall, our *in vitro* ChPO models provided insights into the susceptibility of human ChP epithelial cells to various orthoflaviviruses, which may help explain their clinical neuroinvasiveness.

Beyond demonstrating susceptibility, our demonstration of viral E protein in the ChPO epithelial cells also validated active JEV replication within these cells. In addition, we recovered infectious JEV particles in the CSF fluid within the lumen. Further investigation indicated that the presence of luminal virus was not due to a barrier breach, as permeability assays demonstrated that both mock and infected ChPOs maintained selective barrier integrity, restricting the passage of 3 kDa and 70 kDa dyes. The JEV E monomer alone is approximately 53 kDa; therefore, the intact virion would be significantly larger. Thus, this demonstration of sustained barrier integrity strongly supports a transcellular JEV transport mechanism rather than passive paracellular leakage [49]. This finding mirrors observations in ChPOs infected with the bacterium *Streptococcus suis* [50], suggesting that a nondestructive translocation mechanism is utilized by phylogenetically distinct pathogens. In contrast, previous studies involving viral infections in ChPOs have reported a different outcome, as they specifically showed compromised barrier integrity (as evidenced by tight junction loss and cell death) during SARS–CoV−2 and HSV−1 infections [30, 51]. Nevertheless, our results collectively establish that JEV utilizes a non–destructive transcellular entry mechanism into the lumen, highlighting a distinct strategy for neuroinvasion among pathogens that target the ChP.

The ChP, like other tissues, mounts an intrinsic innate immune response to defend against invading pathogens. Foreign microbes are recognized by pattern recognition receptors (PRRs), a class of receptors that detect and distinguish pathogenic from self–molecules. Components of orthoflavivirus viral particles are sensed by Toll–like receptors (TLRs) and RIG–I–like receptors (RLRs) and cyclic GMP–AMP synthase (cGAS)–STING pathway [52]. In the ChP, the TLRs are the most studied PRRs, particularly regarding the response to bacterial lipopolysaccharide, whereas other types of PRRs have received considerably less attention. Our transcriptomic study detected transcripts for nearly all of these key components, including all TRLs (except *Tlr7* and *Tlr8*), RLRs (*Ddx58*; RIG–I and *Ifih1;* MDA–5), and cGAS–Sting (*Cgas* and *Sting*), and the downstream transcription factors *Irf3* and *Irf7*. Differential expression analysis and independent RT–qPCR validation confirmed the upregulation of these genes following treatment with Poly(I:C) and infection with JEV.

The activation of IRF3/IRF7 and NF–κB leads to the translocation of these transcription factors into the nucleus to promote the production of proinflammatory cytokine genes and two important families of IFNs: type–I and type–III IFNs [53]. Both IFN types are secreted from cells and act in an autocrine or paracrine fashion by binding to their respective cell surface receptors. Type–I IFNs (composed of IFNα, IFNβ, IFNε, IFNκ and IFNω) bind to IFNAR1 and IFNAR2 heterodimers, which are also expressed in all nucleated cells [54, 55]. In contrast, the type–III IFN family (IFNλ1, IFNλ2, and IFNλ3) bind to IFNLR1–3 and IL–10R2 heterodimers, whose expression is largely restricted to epithelial cells [56, 57, 58]. Upon receptor binding, both type–I and III IFNs activate the JAK/STAT signaling cascade, ultimately inducing ISGs, which create an antiviral state in the cells. Our transcriptomic analysis demonstrated upregulation of both IFN types (*Ifnb1* and *Ifnl2/3*), along with key components of the response pathway (*Stat1* and *Irf9*), and numerous ISGs. Our RT–qPCR validation of the tested genes corroborated these findings, except for *Ifnb1*, which remained unchanged throughout the course of infection. Overall, these findings indicate that ChPOs possess functional IFN pathways capable of mounting an antiviral response against JEV infection, similar to those shown in animal models [59].

Although both IFN types are activated by the same PRR pathways and induce largely overlapping sets of ISGs, their spatial responses and kinetics differ. Type–I IFN elicits a universal, rapid, high–magnitude response that declines quickly, whereas type–III IFN responses are restricted primarily to epithelial cells and some immune cells and are slower, lower in magnitude, but generally sustained longer [60]. Interestingly, our RT–qPCR study revealed a universally slower and sustained transcriptional response in selected genes at the late infection timepoint of 96 hpi (Fig. 4C). Moreover, quantification of *Ifnb1* showed no significant upregulation throughout the course of infection, suggesting that either pre–existing IFNβ proteins are sufficient or the ChPO preferentially utilizes other type–I IFN subtypes. One possible alternative could be IFNα, which has been previously demonstrated in the dermal fibroblasts and immune cells of a mouse model to act against Chikungunya virus infection [61]. We also observed a pronounced upregulation of *Ifnl2/3* (approximately 200–fold), which, together with the large number of epithelial cells in the ChPO, suggests that type–III IFN, rather than type–I IFN, may play a dominant role in the response to JEV infection. However, our subsequent investigation using anti– hIFNAR1 and anti–hIFNLR3 indicated that both type–I and type–III IFNs offer protection against JEV infection, although type–I IFN elicits a superior response.

Previous studies have demonstrated that type–III IFNs have protective effects against several viruses in human lung and gut epithelial cells; however, studies on ChP are lacking [62, 63]. Our findings are consistent with studies in which *Ifnar*^-/-^ mice, lacking a type–I IFN response, showed high susceptibility to human HSV–1 and Langat virus, despite having intact type–III IFN responses [19, 64]. Moreover, because our ChPOs contain JEV–susceptible cell types other than epithelial cells, an inhibition of type–III IFN responses in the epithelial cells alone may not be sufficient to increase virus susceptibility if the robust type–I IFN response in those other cell populations remains active.

Understanding the role of IFN responses at the brain barrier is crucial, as the results from the present study suggest that the IFN response might represent a crucial mechanism for protecting against JEV neuroinvasion at the blood–CSF barrier. A recent study using spatial transcriptomics in JEV–infected mouse brain showed that ISGs were highly expressed in ChP as soon as day 1 after footpad inoculation, when the virus was not present in the brain, and they were expressed at even higher levels when viral RNA could be detected in the brain on day 4 after infection [59]. Furthermore, a growing body of evidence suggests that autoantibodies that neutralize IFN responses might be associated with increased severity in several infectious diseases, including SARS–CoV–2, MERS–CoV, influenza, and HIV–1 [65, 66, 67, 68]. Alarmingly, these autoantibodies have been detected in 40% of patients with WNV encephalitis, suggesting that this population could be at a greater risk of viral neuroinvasion [69]. In contrast, an excessive IFN response could have a negative impact on infected patients, as indicated by cognitive impairments occurring in SARS–CoV–2 infected animal models and patients [70, 71]. Therefore, further in–depth studies are critically needed to advance our understanding of how the ChP orchestrates a response against invading viruses. The use of state–of–the–art technologies, such as single–cell RNA sequencing or spatial RNA sequencing, will help generate a more precise understanding of the interplay between different types of IFN responses and the various cell populations within the ChPOs.

The present study has a major limitation, which the organoid model inherently lacks immune cells. *In vivo,* the ChP harbors a unique population of microglia and serves as a crucial gateway for infiltration of peripheral immune cells, such as T cells, B cells, macrophages, and dendritic cells, into the CNS [72, 73]. These immune components are vital for immunosurveillance and for preventing pathogen invasion. However, the presence of immune cells can be a double–edged sword as the cells can contribute to neuroinflammation and potentially cause damage to the brain barriers and the CNS. Notably, we detected the upregulation of genes encoding chemokines (such as *Cxcl10, Cxcl11, and Ccr5*), which are known to recruit immune cells and enhance neuroinflammation [74, 75, 76]. Therefore, future studies should incorporate immune components, such as co–cultures of iPSC–derived microglia or trans–well systems containing peripheral immune cells, to generate a more accurate model of the complex interplay between the ChP and the immune system, which appears to modulate viral infection and barrier integrity [30, 77].

## Conclusion

The ChP, which forms the blood–CSF barrier, is recognized as a critical defense against neurotropic pathogens, yet the precise mechanism by which neuroinvasive orthoflaviviruses bypass this barrier remains largely unknown. We addressed this knowledge gap by establishing *in vitro* ChPOs derived from CD34^+^–iPSCs that successfully recapitulated characteristics of the blood–CSF barrier, including CSF–like fluid secretion and selective barrier integrity. We demonstrated that our ChPOs were susceptible to various neurotropic orthoflaviviruses (JEV, ZIKV, and DENV2). Notably, despite demonstrating productive JEV replication within epithelial cells and the release of infectious viruses into the CSF–like fluid, we observed no compromise or breach of the epithelial barrier integrity. This finding strongly supports a transcellular neuroinvasion mechanism whereby JEV accesses the ChPO lumen by infecting the ChP epithelial cells, which subsequently release viral particles apically without causing damage to the barrier. Furthermore, our ChPOs exhibited an intrinsic innate immune response to JEV, as evidenced by the significant upregulation of the IFN induction and response pathways. Our functional studies revealed that both type–I and type–III IFNs play an important role in controlling JEV infection, with type–I IFNs being superior. Collectively, this ChPO model provided a valuable *in vitro* platform for studying mechanisms of viral infection and the interaction of the human ChP. The findings revealed the JEV entry route as a nondestructive transcellular neuroinvasion and determined the intrinsic response to JEV infection, thereby advancing our understanding of ChP barrier responses to orthoflavivirus infection.

## Declarations

### Ethical approval and consent to participate

The use of human iPSC clone MURAi003-A, derived from CD34+ hematopoietic stem cells from a healthy Thai donor (permit number MURA2022/321), followed the guidelines and regulations approved by the Ethical Committee, Faculty of Ramathibodi Hospital, in accordance with the Declaration of Helsinki and the International Organizations of Medical Sciences (CIOMS) Guidelines 2016. Infectious work was performed in the certified Biosafety Level 2 lab, following the guidelines and regulations approved by the Ramathibodi Hospital Institutional Biosafety Committee (permit number RAMA-IBC 2023-001). Consent to participate declaration is not applicable.

### Consent for publication

Not applicable

### Availability of data and materials

The raw RNA–sequencing data generated for this study have been deposited in the European Nucleotide Archive (ENA) at EMBL–EBI under accession number PRJEB102258. All data generated or analyzed during this study are included in this published article and its supplementary information files.

### Competing interests

The authors declare that they have no competing interests

### Funding

This work was supported by funding from the Faculty of Medicine Ramathibodi Hospital, Mahidol University, and Mahidol University Strategic Research Fund: 2024 to NC. NJ was supported by the Thailand Program Management Unit for Human Resources & Institutional Development, Research and Innovation (PMU–B), NXPO, grant number B05F640142.

### Authors’ contributions

NC, YK, and NA conceptualized the main research questions and experimental approach. NC, YK, AB, NA, and NJ designed the experiment. UA, CM, PK, and BS provided guidance and experimental design on specific techniques. YK, AB, PP, CP, and NC performed and analyzed the experiment and contributed to the analysis and discussion of the results. NC, YK, and AB wrote the first draft of the manuscript with extensive revision and advice from NA and NJ. All authors revised and approved the final version of the manuscript.

## Acknowledgements

We thank Dr. Kran Suknuntha and the Anatomical Pathology Laboratory, Chakri Naruebodindra Medical Institute (CNMI), Faculty of Medicine Ramathibodi Hospital, for facilitating cryosection and tissue staining. We are also grateful to Dr. Kwanchanok Uppakara for training in the confocal imaging facility at the Medical School Ramathibodi, Faculty of Medicine Ramathibodi Hospital. We thank Dr. Tanapat Pornsukjantra, Faculty of Medicine Ramathibodi Hospital, for iPSC culture training, and Dr. Suparerk Borwornpinyo, Faculty of Science, Mahidol University, for allowing us to use their organoid culture facility. Lastly, we also gratefully thank Dr. Pimonrat Ketsawatsomkron and Dr. Kenjiro Muta from the Faculty of Medicine Ramathibodi Hospital for providing comments on the project. We thank Monica Madore from Scribendi for editing a draft of this manuscript. Schematics in figures 3-5 were created with BioRender.com.

## Supplementary file 1

Supplementary table 1 and figure 1

## Supplementary file 2

DEGs of ChPOs treated with Poly(I:C) compared to mock. RNA sequencing was performed on the ChPOs treated with 20 µg/mL Poly(I:C) for 24 h and mock samples (3 organoids/sample; 3 samples per condition). The gene list was selected from the transcript with |log_2_ fold change| ≥ 1 and adjusted-*p* value < 0.05.

## Supplementary file 3

DEGs of ChPOs infected JEV compared to mock. RNA sequencing was performed on the ChPOs infected with 10^5^ PFU JEV strain Nakayama for 24 h and mock samples (3 organoids/sample; 3 samples per condition). The gene list was selected from the transcript with (|log_2_ fold change| ≥ 1) and adjusted-*p* value < 0.05.

## Notes

### Competing Interest Statement

The authors have declared no competing interest.

