## Supplementary file 1 for "Mechanism of neuroinvasion and interferon response in human choroid plexus organoids during Japanese encephalitis virus infection"

**Table S1. Primers**

| Genes | Primers | Sources |
|---|---|---|
| *Bax* | F – 5’ AGTAACATGGAGCTGCAGAGGAT 3’<br>R – 5’ GCTGCCACTCGGAAAAAGAC 3’ | [1] |
| *Ddx58* | F – 5’ AGACCCTGGACCCTACCTACA 3’<br>R – 5’ CTCCATTGGGCCCTTGTTGT 3’ | [2] |
| *Gapdh* | F – 5’ ATGACATCAAGAAGGTGGTG 3’<br>R – 5’ CATACCAGGAAATGAGCTTG 3’ | [3] |
| *Ifih1* | F – 5’ ATTCAGGCACCATGGGAAGTG 3’<br>R – 5’ TTTGGTAAGGCCTGAGCTGGA 3’ | [2] |
| *Ifit1* | F – 5’ AGAAGCAGGCAATCACAGAAAA 3’<br>R – 5’ CTGAAACCGACCATAGTGGAAAT 3’ | [4] |
| *Ifnb1* | F – 5’ GCCGCATTGACCATCTAT 3’<br>R – 5’ GTCTCATTCCAGCCAGTG 3’ | [5] |
| *Ifnl2/3* | F – 5’ GCCACATAGCCCAGTTCAAG 3’<br>R – 5’ TGGGAGAGGATATGGTGCAG 3’ | [5] |
| *Irf3* | F – 5’ CCAAGAGGAAGTCATGTG 3’<br>R – 5’ TAGCCTGGAACTGTGTAG 3’ | [6] |
| *Nme5* | F – 5’ GACCTAAGGAATGCACTTCATGG 3’<br>R – 5’ GTGAGTCCTTCAAGCAGAGTTGG 3’ | Origene, NM_003551 |
| *Ocln* | F – 5’ TCAGGGAATATCCACCTATCACTTCAG 3’<br>R – 5’ CATCAGCAGCAGCCATGTACTCTTCAC 3’ | [7] |

| **Genes** | **Primers** | **Sources** |
|---|---|---|
| *Tlr3* | F – 5’ GAGGCGGGTGTTTTTGAACTAGAA 3’<br>R – 5’ AAGTCAATTGTCAAAAATAGGCCT 3’ | [8] |
| *Usp18* | F – 5’ CCTGAGGCAAATCTGTCAGTC 3’<br>R – 5’ CGAACACCTGAATCAAGGAGTTA 3’ | [4] |
| *Zo1* | F – 5’ GTGTTGTGGATACCTTGT 3’<br>R – 5’ GATGATGCCTCGTTCTAC 3’ | [7] |

**Supplementary figure**

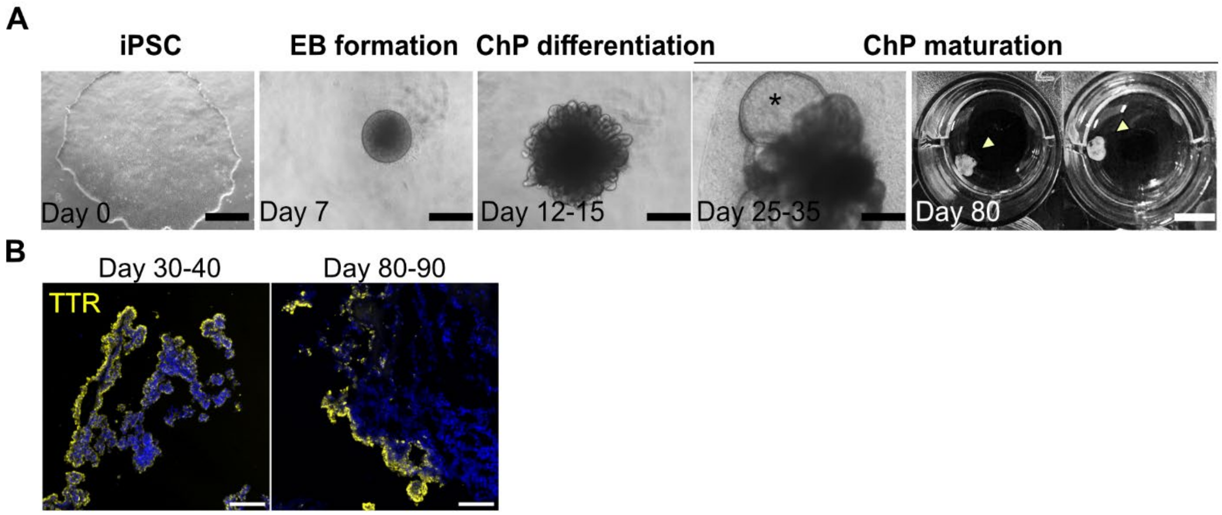


**Fig S1. Generation and characterization of ChPO derived from BYS0113 (ATCC).** (A) Representative images illustrating the generation and maturation of BYS0113 choroid plexus organoids (ChPOs). The observed differentiation timeline and process were consistent with those previously observed in the MURAi003-A-derived ChPOs. Black scale bars = 500 μm and white scale bar = 1 cm. **(B)** Immunofluorescence staining of BYS0113-derived ChPOs with TTR (yellow) and DAPI (blue). Scale bars = 100 μm.
